# A Purkinje cell model that simulates complex spikes

**DOI:** 10.1101/2020.05.18.102236

**Authors:** Amelia Burroughs, Nadia L. Cerminara, Richard Apps, Conor Houghton

## Abstract

Purkinje cells are the principal neurons of the cerebellar cortex. One of their distinguishing features is that they fire two distinct types of action potential, called simple and complex spikes, which interact with one another. Simple spikes are stereotypical action potentials that are elicited at high, but variable, rates (0 – 100 Hz) and have a consistent waveform. Complex spikes are composed of an initial action potential followed by a burst of lower amplitude spikelets. Complex spikes occur at comparatively low rates (~ 1 Hz) and have a variable waveform. Although they are critical to cerebellar operation a simple model that describes the complex spike waveform is lacking. Here, a novel single-compartment model of Purkinje cell electrodynamics is presented. The simpler version of this model, with two active conductances and a leak channel, can simulate the features typical of complex spikes recorded *in vitro*. If calcium dynamics are also included, the model can capture the pause in simple spike activity that occurs after complex spike events. Together, these models provide an insight into the mechanisms behind complex spike spikelet generation, waveform variability and their interactions with simple spike activity.

## 1 Introduction

Purkinje cells fire two distinct types of action potential, called complex spikes and simple spikes, in response to two different types of excitatory input relayed via climbing fibre and mossy fibre afferents, respectively (Palay and Chan-Palay, 1974; Ito, 1984; Eccles, 2013). The mossy fibres connect indirectly to the Purkinje cells via granule cells; these, in turn signal to the Purkinje cells along the parallel fibres. Activity within the mossy fibre - granule cell - parallel fibre pathway modulates the already intrinsic simple spike activity of Purkinje cells, which averages at ~ 40 Hz for spontaneous activity in rats (Armstrong and Rawson, 1979), cats (Thach, 1967b) and monkeys (Fu et al., 1997) but can range from 0 – 200Hz (Chen et al., 2016). In marked contrast, complex spikes are generated at only ~ 1 Hz (Lang et al., 1999) in response to activity within the direct climbing fibre pathway and are characterised by a prolonged, multi-peaked action potential (Campbell and Hesslow, 1986). The initial spike in a complex spike has a larger amplitude to the subsequent peaks, these smaller peaks are known as spikelets, which are elicited at ~ 600Hz (Warnaar et al., 2015; Burroughs et al., 2017). The complex spike is followed by an extended refractory period (Voogd and Glickstein, 1998; Eccles et al., 1966; Fujita, 1968).

Complex spikes, and their interactions with simple spikes, are central to theories of cerebellar function (Campbell and Hesslow, 1986; Eccles, 2013; Ito, 1984, 2011; Yang and Lisberger, 2014). While most theories consider complex spikes as unitary events, there is increasing evidence to suggest that changes in the complex spike waveform plays an important role. For example, there is recent evidence that behaviourally salient information is transmitted by variations in complex spike waveform and the duration of complex spikes differs between spontaneous and sensory-evoked events (Maruta et al., 2007; Najafi and Medina, 2013). Complex spike duration also appears to affect plasticity: the magnitude of motor learning and synaptic plasticity in awake monkeys correlates with complex spike duration (Yang and Lisberger, 2014). It has also been proposed in Houghton (2014) that the refractory period which follows a complex spike has a role in plasticity at the granule cell - Purkinje cell synapse.

There is also emerging evidence for an interaction between complex spike waveform changes, simple spike activity and behaviour (Yang and Lisberger, 2014; Streng et al., 2017). The best known interaction is the pause in simple spike activity that follows a complex spike, but an increasing number of studies indicate a relationship between simple spike firing dynamics and complex spike activity (Mano, 1970; Gilbert, 1976; Campbell and Hesslow, 1986; Hashimoto and Kano, 1998; Servais et al., 2004; Maruta et al., 2007; Warnaar et al., 2015; Burroughs et al., 2017). Variations in complex spike waveform may dynamically regulate the simple spike response of Purkinje cells in advance of changes in behaviour in order to determine the motor outcome (Streng et al., 2017). Furthermore, changes in the waveform of the complex spike may serve a homeostatic role in maintaining Purkinje cell simple spike activity within a useful operational range (Burroughs et al., 2017).

Purkinje cell responses are highly variable: there is considerable variation in firing rates, input response properties, morphology and complex spike waveform. There are, however, some features that appear to be generic to Purkinje cells: the spikelets that make up the complex spike have a significantly smaller amplitude than the initial spikes and there is an extended refractory period after a complex spike. The purpose of this paper is to suggest mechanisms to support these features by describing simple models that can simulate them.

Given the evidence that a dynamic interplay between complex spike waveform and simple spike activity underlies cerebellar operation, there is a need to gain a mechanistic understanding of the generation of the complex spike and its interaction with simple spiking. Despite the existence of a range of computational models to address various features of Purkinje cells, a simple model that captures complex spike generation, waveform variability and interactions with simple spike activity is lacking. In the current paper we explore through simulation the hypothesis that a restricted number of channel-dynamics can explain complex spike waveform. Two models are presented: a three-current model that shows that three channels, a leak channel and two active channels, are sufficient to simulate the complex spike waveform and a five-current model, which demonstrates how some of the interactions between simple spikes and complex spikes can be explained by calcium dynamics.

These models serve as a description of the essential ion-channel dynamics supporting complex spike production and should enable further analysis of complex spiking and its relationship to simple spiking. In this way, the models complement the large and detailed models described by Veys et al. (2013); Zang et al. (2018) and give a starting point for a model which also incorporates the interaction between simple spike rate and complex spike waveform described in Burroughs et al. (2017).

## 2 Methods

In this paper two single-compartment models of the Purkinje cell somatic voltage dynamics are presented. The three-current model has a leak channel and two voltage-gated channels:

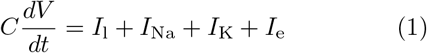

where *C* =1 *μ*F /cm^2^ is the membrane capacitance, *I_l_* is a leak current,

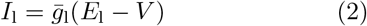

with 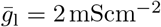 and *E_l_* = −88 mV. *I*_e_ stands for the external input current. This is described below and is made up of a background current and a synaptic current entering the soma from the dendrites. *I*_Na_ and *I*_K_ stand for the sodium current and the potassium current; the sodium current includes the resurgent dynamics that are typical of Purkinje cells (Raman and Bean, 1997, 2001; Khaliq et al., 2003; Khaliq and Raman, 2006). Since complex spikes are principally sodium spikes (Stuart and Häusser, 1994), this model contains the least possible number of currents; the key contribution of this model is to demonstrate that the sodium resurgence found in Purkinje cells is sufficient as a mechanism for producing the complex spike waveform.

The second model includes some of the effects of calcium dynamics and will be referred to as the five-current model. It has

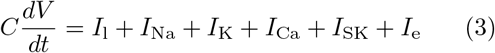

In addition to the three-current model described above in Eq 1, the five-current model also has a calcium channel, with current *I*_Ca_ and a calciumgated potassium channel, with current *I_SK_*. These two additional currents are also described below; they were selected because they are a prominent channel type in Purkinje cells and the calcium dynamics has timescales in the range required to produce the refractory period after a complex spike. Purkinje cell channels that are not included in either model are described in the Discussion.

### 2.1 Input currents

In the Purkinje cell dendrites, climbing fibre activation evokes a sustained depolarising calcium transient (Kitamura and Häusser, 2011). This transient can include spikes and the number of action potentials in the presynaptic climbing fibre burst can influence the number of these dendritic calcium spikes (Llinas and Sugimori, 1980a; Kitamura and Häusser, 2011; Davie et al., 2008; Mathy et al., 2009). However these calcium spikes are not well propagated from the dendrites into the soma (Davie et al., 2008) and as such the Purkinje cell soma most likely receives the dendritic climbing fibre signal as a depolarisation plateau carried mostly by calcium and non-inactivating, passive sodium channels (Llinas and Sugimori, 1980b; Knäpfel et al., 1991; Llinas and Nicholson, 1971; Stuart and Häusser, 1994). This depolarisation plateau drives somatic complex spike formation, which is initiated in the proximal axon (Stuart and Häusser, 1994; Davie et al., 2008; Palmer et al., 2010).

The current that the Purkinje cell soma receives following climbing fibre activation from the dendrite is modelled as the difference of two decaying exponentials and fitted to mimic the synaptic input given in Davie et al. (2008). For a complex spike at *t* = 0 this is:

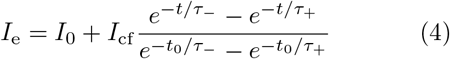

where to is the time the double exponential function reaches its maximum:

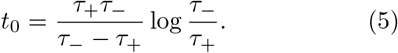

*I*_0_ = 62 *μ*Acm^-2^ and *I*_0_ = 67 *μ*Acm^-2^ are background currents in the three-current model and five-current model respectively, *I*_cf_ = 52 μAcm^-2^ and *I*_cf_ = 71 *μ*Acm^-2^ are the maximum amplitudes of the climbing fibre current in the three-current model and the five-current model respectively, *τ*_+_ = 0.3 ms and *τ*_−_ =4 ms are the rise and decay timescales respectively and are the same for both the three-current and five-current model.

### 2.2 Ion channels

The current through the sodium channel was modelled using a Markovian scheme developed by Raman and Bean (2001), see Table 1. This scheme models the dynamics of both the transient and resurgent gates. The current is

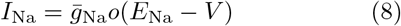

*o* is the fraction of gates in the open state *O*, as described in Table 1 and 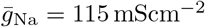.

**Table 1:**
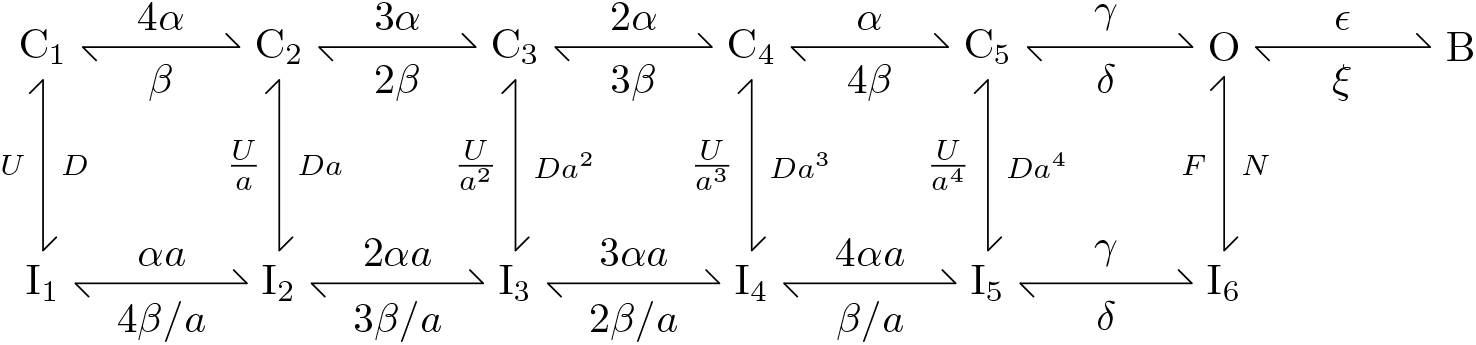
Current through the resurgent sodium channel is described using a Markovian Scheme. *C*_1_-*C*_5_ describe sequential closed configurations; *I*_1_-*I*_6_ describe sequential inactivated states; O represents the open channel configuration, sodium ions will only pass through the channel when it is in this state. *B* describes a second inactivated state where the channel is likened to being in an open-but-blocked configuration that is non-conducting. Return from this state back to the closed, or inactivated, states must occur through the open (*O*) configuration. The rate constants describing transitions between states; all rate values have been taken from Raman and Bean (2001).

The potassium current has a fast potassium channel of the K_v_3.3 subtype and is modelled as in Masoli et al. (2015):

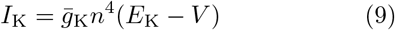

where

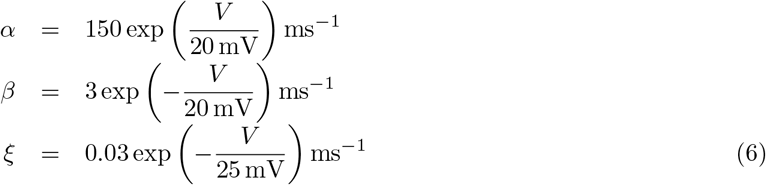

and *γ* = 150ms^-1^, *δ* = 40ms^-1^, *ϵ* = 1.75ms^-1^, *D* = 0.005ms^-1^, *U* = 0.5ms^-1^, *N* = 0.75ms^-1^, *F* = 0.005 ms^-1^ with

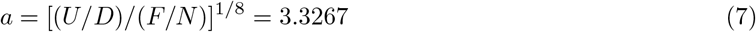

where 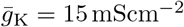 and *E*_K_ = −88mV. Channel kinetics are described with the usual Hodgkin-Huxley channel equation but with:

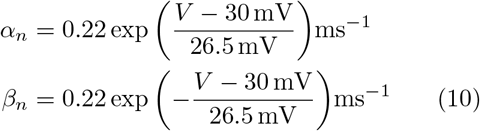

The five-current model extends the three-current model to include calcium. It adds a calcium channel, a model of calcium concentration dynamics in the soma and a calcium-gated potassium channel. The calcium channel is a P/Q type calcium channel with equations and parameter values the same as those in

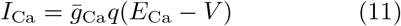

with 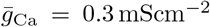 and *E*_Ca_ = 135 mV. *q* is the proportion of gates in the open state. Channel kinetics are given by

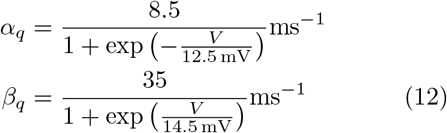

The intracellular calcium concentration dynamics are taken from Sterratt et al. (2012); Koch and Segev (1998)

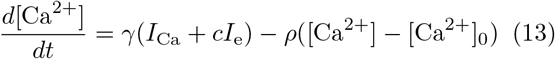

Thus, calcium enters the soma as part of the *I*_e_ input current and through calcium channels; *c* = 0.02 is a measure of what fraction of Ie is composed of calcium and *γ* is a conversion factor which converts the current to a calcium concentration flux. As well as quantifying the charge of each ion, this factor needs to account for the distribution of calcium in the soma; unlike current which depends on membrane area, concentration depends on volume. However, in the soma the calcium is sequestered in a thin layer near the membrane. Hence, *γ* = *0.01/zFδr* cm^-2^ where *z* = 2 is the electrical charge of the calcium ion, *δ*r = 0.2 *μ*m is the width of the somatic submembrane shell and F is Faraday’s constant. In the absence of calcium inflow the calcium concentration returns with rate constant *ρ* = 0.02ms^-1^ to a baseline value [Ca^2+^]_0_ = 30nM cm^-3^ through intracellular buffering (Fierro et al., 1998; Airaksinen et al., 1997); this value has been chosen so that the timecourse matched the description in Eilers et al. (1995) of calcium concentration dynamics in the subsomatic cell of Purkinje cells.

The calcium activated potassium channel is an SK, small conductance, channel, (Lancaster et al., 1991). The channel dynamics are taken from Gillies and Willshaw (2006) and are described by

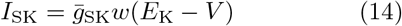

where 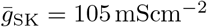. Channel kinetics are

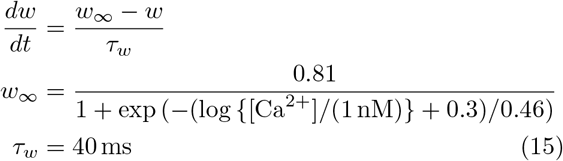

### 2.3 Implementation and model fitting

Parameter values are given in Table 2. Most parameter values used in this modelling study were taken from experimental studies. However, some parameters that are not easily measured experimentally are optimised here so that the complex spike waveform shape had the best fit to complex spikes recorded experimentally *in vitro*, see Figure 1D in Davie et al. (2008).

**Table 2:** Parameter values and their associated references. The parameters marks with a star (*) have been handtuned, where there is a reference it is to the starting value given beside the star. The parameters marked with a double dagger (‡) have been chosen to fit the dynamics of the somatic voltage change recorded in response to climbing fibre activity in Davie et al. (2008).

| Parameter | Value | Reference |
| --- | --- | --- |
| $C$ | $1 \mu\text{F cm}^{-2}$ | (Roth and Häusser, 2001) |
| $\bar{g}_l$ | $2 \text{ mScm}^{-2}$ | (Rapp et al., 1994) |
| $\bar{g}_{\text{Na}}$ | $47.2 \text{ mScm}^{-2}$ | $\star \dagger$ |
| $\bar{g}_{\text{K}}$ | $200 \text{ mScm}^{-2}$ | $\star 198 \text{ mScm}^{-2}$ (Akemann and Knopfel, 2006) |
| $\bar{g}_{\text{Ca}}$ | $0.3 \text{ mScm}^{-2}$ | $\star 0.5 \text{ mScm}^{-2}$ (Miyasho et al., 2001) |
| $\bar{g}_{\text{SK}}$ | $78 \text{ mScm}^{-2}$ | $\star 120 \text{ mScm}^{-2}$ (Rubin and Cleland, 2006) |
| $E_l$ | $-88 \text{ mV}$ | (Masoli et al., 2015) |
| $E_{\text{K}}$ | $-88 \text{ mV}$ | (Masoli et al., 2015) |
| $E_{\text{Na}}$ | $45 \text{ mV}$ | (De Schutter and Bower, 1994a) |
| $E_{\text{Ca}}$ | $135 \text{ mV}$ | (De Schutter and Bower, 1994a) |
| $[\text{Ca}^{2+}]$ | $30 nM \text{ cm}^{-3}$ | (Kano et al., 1995) |
| $\delta r$ | $0.2 \mu\text{m}$ | |
| $I_0$ | $62.46 \mu\text{A cm}^{-2}$ | $\star$ |
| $I_{\text{cf}}$ | $90 \mu\text{A cm}^{-2}$ | ‡ (Stuart and Häusser, 1994) |
| $\tau_+$ | $0.3 \text{ ms}$ | ‡ (Stuart and Häusser, 1994) |
| $\tau_-$ | $4 \text{ ms}$ | ‡ (Stuart and Häusser, 1994) |

**Figure 1:**
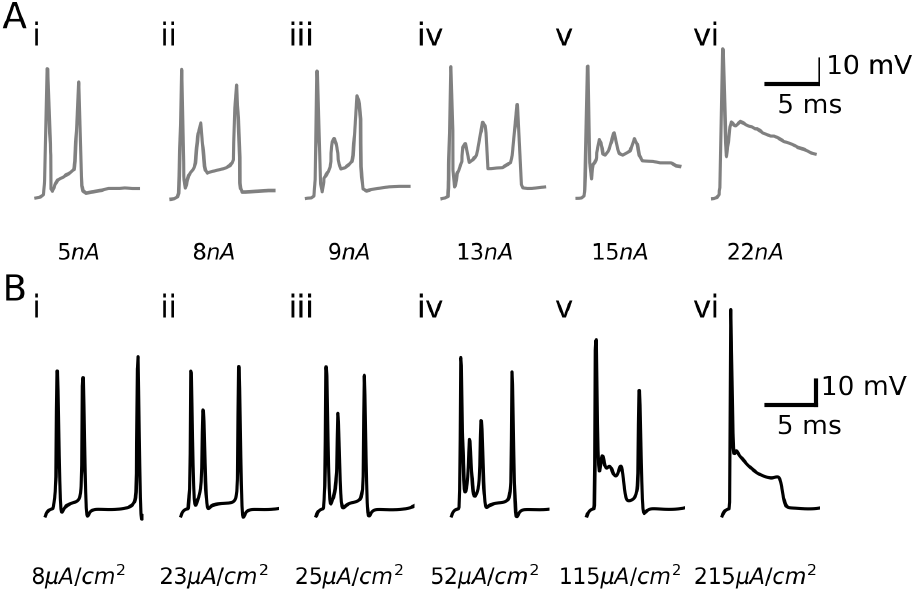
Simulated complex spike responses to increasing amplitudes of injected current. The top row **(A)** shows in *vitro* Purkinje cell recordings from Davie et al. (2008) in response to increasing (from 5 *nA* **(Ai)** through to 22 *nA* **(Avi)**) injections of climbing-fibre-like synaptic current. The bottom row **(B)** shows the result of the three-current model simulation in response to increasing injections of climbing-fibre-like somatic current (from 8 *uA/cm2* **(Bi)** through to 215 *uA/cm2* **(Bvi)**). The structure of the complex spikes, and the manner of their response to increasing amplitudes of injected current, are very similar when comparing the *in vitro* responses and the output of our model. The models presented here were fitted to the complex spike recorded in **(Aiv)**.

Simulations were run in Python 2.7, OS on a CPU. Models were simulated as ordinary differential equations and integration was performed explicitly using the `scipy.integrate.ode` package using the `BDF` option suitable for stiff problems, this implements a backward differentiation formula with a Jacobian estimated by finite differences (Byrne and Hindmarsh, 1975). The timestep, *dt*, was chosen as 0.0025 ms. One second of Purkinje cell activity can be simulated in 1 minute 20 seconds.

Optimization was performed using a mixture of error minimization and hand-tuning techniques. In order to have a definite target in tuning parameters the model was optimized to simulate features of a typical complex spike recorded *in vitro* in Davie et al. (2008) and illustrated here, with permission, in Fig. 1Aiv. This complex spike was chosen because complex spikes with three spikelets are most common (Burroughs et al., 2017). Although this is a specific experimental result; it exhibits those general features of complex spikes we wanted to simulate in our model and so it serves as a useful target for parameter fitting. Specifically, the model was fitted to specific features of the complex spike waveform that included the timing of individual spikelets within the complex spike, the order of spikelet amplitudes within the complex spike and overall complex spike duration.

The error used was the squared distance between the simulated Purkinje cell voltage trace in response to climbing fibre input and the typical complex spike recorded *in vitro*, Fig. 1Aiv. Error minimization used the downhill simplex algorithm^1^. The square-error can be unstable for spike-like profiles, given that minor changes in the timing of individual spikelet peaks can generate huge changes in the square error even if other spike features are well described. This could be addressed through a time rescaling; here, though, the error minimization was combined with hand-tuning.

We hand-tuned by running simulations thousands of times with different combinations of physiologically plausible parameter values to create large grids of simulated complex spike images and then selecting the best by eye. Current injections necessary for simulating complex spikes were comparable to those necessary to reproduce the same complex spike *in vitro*. In the model, injections of climbing-fibre-like somatic currents varied from 8 μA/cm^2^ through to 215 μA/cm^2^ Fig. 1. Assuming a surface area of 500 μm^2^ this would correspond to current injections between 0.4 nA and 10.75 nA.

## 3 Results

The three-current and five-current model were tested. After fitting those parameters that are not known from experiment, the three-current model simulates complex spikes whose structure exhibits those features which are broadly typical in Purkinje cells; the five-current model in addition, simulated the extended refractory period which follows complex spikes. Table 2 gives a list of the model parameters, giving the source for parameters that have been fixed to an experimental value and noting which parameters were adjusted to fit the target complex spike.

The three-current model includes a leak channel, a sodium channel with both transient and resurgent sodium gates, and a fast potassium channel. The main features of the complex spike shape are found in this three-current model, Fig. 1. With increasing injections of climbing-fibre-induced somatic current, the number of spikelets in the simulated complex spike increased from one to three. Each additional spikelet had successively greater amplitudes than the spikelet before it. For complex spikes with only two spikelets the latency from the first to the second spikelet shortens with increasing current. As the current is increased the number of spikelets increases to three. Further increases cause the amplitude of the spikelets to decrease, before the spikelets are lost altogether and replaced by a depolarisation plateau. The model was robust to injections of greater parallel fibre current and the subsequent increases in the rate of simple spike firing. To indicate the relationship between the complex spike and the input current these are plotted together in Fig. 2 for the specific complex spike seen in Fig. 1Biv.

**Figure 2:**
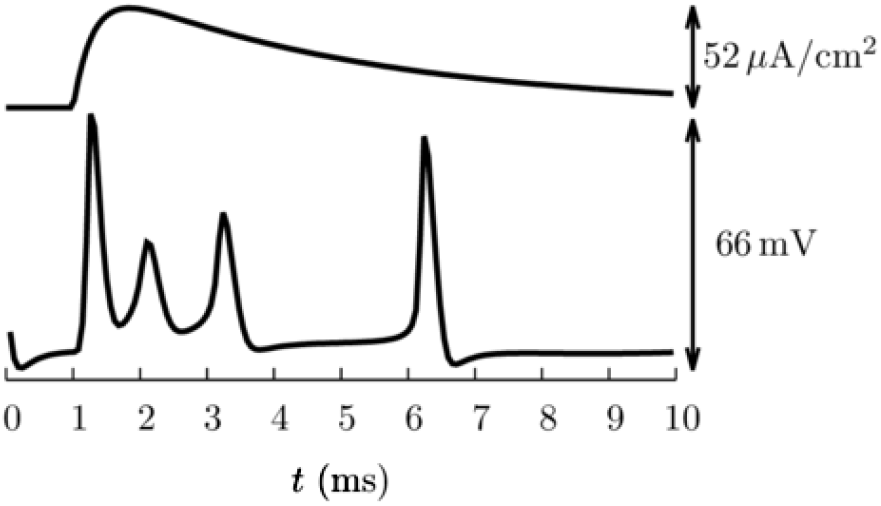
Comparison of the input current and complex spike. This shows 10 ms of the **(above)** input to the three-current model and **(below)** complex spike response; the complex spike is the same one shown in Fig. 1Biv. The input departs from *I*_o_ at *t* =1 ms.

The contributions of each of the individual currents are outlined in Fig. 3. The interplay between the resurgent sodium and fast potassium currents is critical in generating the spikelets in the complex spike. Spikelet formation is absent following the removal of the K_v_3.3 current, in line with experimental investigation (Zagha et al., 2010), but there is still a depolarisation plateau following the injection of climbing fibre current. In the absence of the sodium current, spikelets are abolished, as are simple spikes, and the resurgent sodium current is apparent after the initial spike of the complex spike, but spikelet generation fails. The combination of active sodium and potassium currents is therefore necessary for modelling complex spikes.

**Figure 3:**
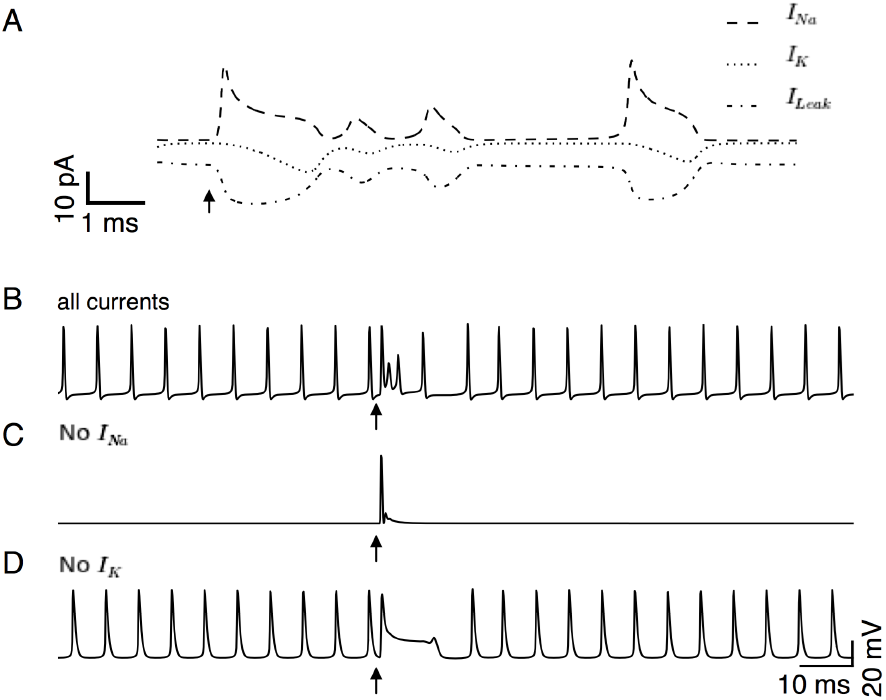
Currents involved in complex spike formation. The contribution of each current in the three-current model (*I*_Na_, dash; *I*_K_, dot; *I*_l_, dashdot) to complex spike generation, **(A)**. **(B)** shows the voltage trace of the simulated Purkinje cell in the three-current model. The resulting simulated Purkinje cell activity following removal of each of the active conductances (**(C)**: *I*_Na_, **(D)**: *I*_K_). Arrows indicate the times of the simulated climbing-fibre-like current injections.

It is possible to increase the number of spikelets generated in the complex spike waveform to more than three by increasing the decay time constant *τ_* in the injected current I_e_, Fig. 4. This is in line with experimental studies, which have shown that an increase in the number of bursts in the pre-synaptic olivary action potential generates an increase in dendritic calcium spikes, which is in turn linearly related to the number of spikelets in, and duration of, somatic complex spikes (Mathy et al., 2009). This is presumably due to an increase in the duration of the climbing-fibre-induced somatic depolarisation. The injected current, modelling the climbing-fibre-like input to the soma *in vitro* in Davie et al. (2008), has a difference-of-exponentials form with a rise time *τ*_+_ = 0.3 ms and decay time *τ* − = 4 ms. When the decay time *τ*_−_ is increased from 4 ms to 18 ms the number of spikelets increases. Spikelet amplitudes are also affected. In comparison to complex spikes simulated with decay time *τ*_−_ = 4 ms, when the rise time τ+ is larger, spikelets in the middle of simulated complex spikes were reduced in amplitude but the amplitude of those towards the end of the complex spike are increased. This is strikingly similar to the Purkinje cell responses described experimentally *in vitro* in Monsivais et al. (2005).

**Figure 4:**
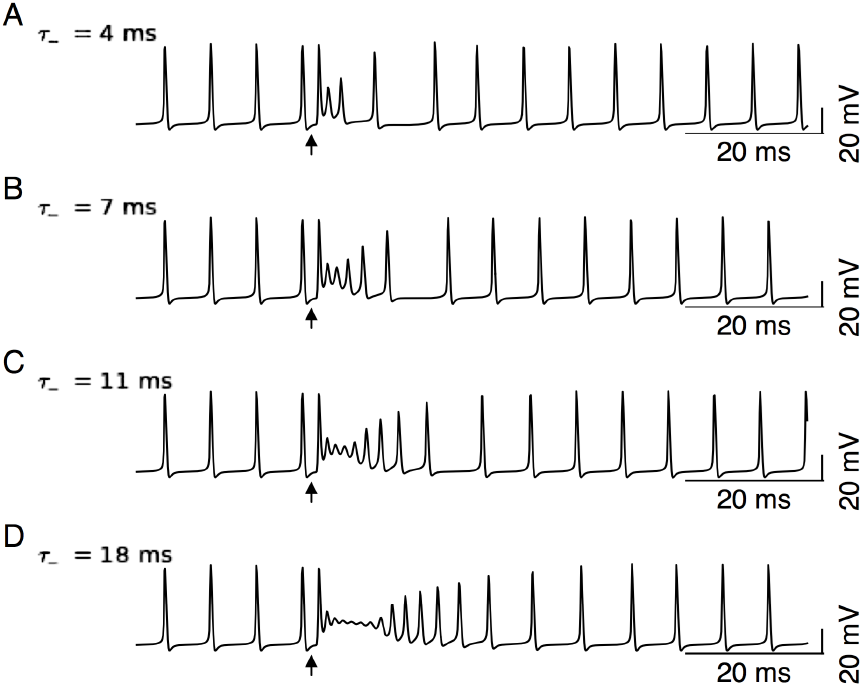
Response of simulated complex spikes to increasing durations of injected current. The time constant describing the decay of the climbing-fibre-like current injection affected the shape of the complex spike waveform. Increasing *τ*_−_ from 4 ms **(A)**, 7 ms **(B)**, 11 ms **(C)** and 18 ms **(D)** increases the number, but decreases the amplitude, of secondary spikelets within the complex spike. Arrows indicate the times of the simulated climbing-fibre current injections. The shape of these simulated complex spikes are similar to those recorded in Purkinje cells *in vitro*, see Figure 6B in Monsivais et al. (2005).

In addition to generating complex spikes, the three-current model also produces simple spikes, Fig. 5A. A distinctive feature of Purkinje cell activity is the pause in the ongoing simple spike activity following a complex spike event (Bell and Grimm, 1969; Granit and Phillips, 1956; Thach, 1967a), which typically lasts for 50 ms. In the three-current model there is only a very brief cessation in the ongoing simple spike train following the complex spike event, a pause that was no greater than the ≈ 15 ms time between two simple spikes, Fig. 5A. Thus, the three-current model does not simulate the interaction between complex spikes and simple spikes. To address this, two further channels were added: a voltage-gated calcium channel and a calcium-gated potassium channel.

**Figure 5:**
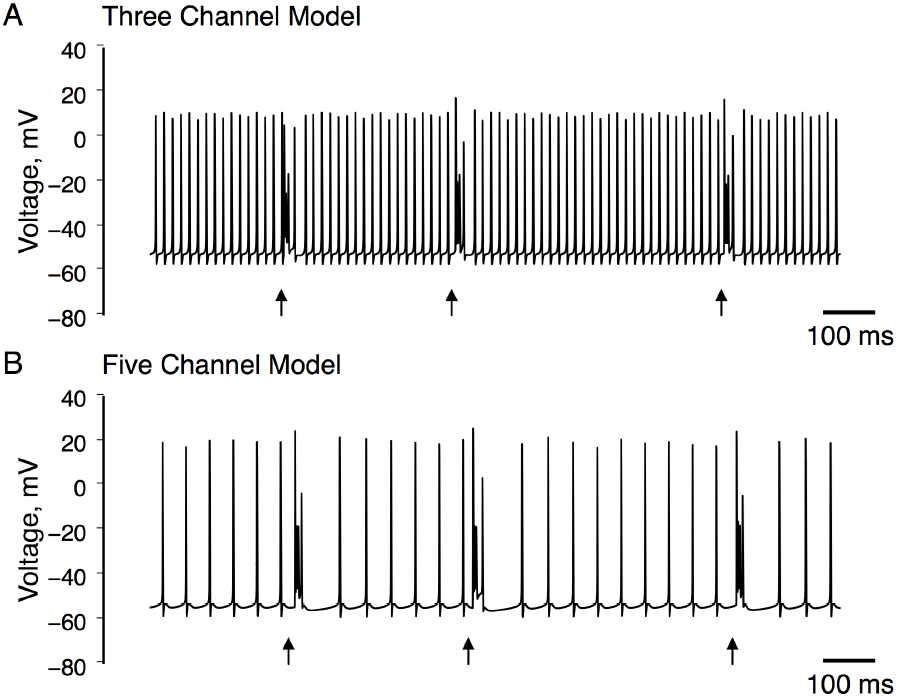
Longer time-base Purkinje cell simulations showing complex spike and simple spike activity. Two second simulation of the three-current model **(A)** and the five-current model **(B)**. Arrows indicate the times of climbing-fibre-like current injections.

The baseline simple spike rate in the five-current model is ~ 40 Hz, Fig. 5B. This is in line with the rate of spontaneous Purkinje cell simple spike activity recorded *in vivo* in anaesthetized rat, see for example Burroughs et al. (2017). The ongoing simple spike activity was interrupted by simulated complex spikes generated in response to injections of climbing-fibre-like current. Following individual complex spikes there was a transient cessation in the simple spike train that resembles Purkinje cell behaviour observed in *vitro* and *in vivo*. This pause lasted for ~ 40 ms before returning to baseline levels of simple spike activity. This pause was greater than expected if complex spikes and simple spikes are independent (simple spike interespike interval divided by two, see Xiao et al. (2014)), unlike in the three-current model. Simulated complex spikes in the five-current model, in response to increasing time constants describing the decay of the injected climbing fibre input, behave similarly to those simulated in the three-current model, Fig. 6.

**Figure 6:**
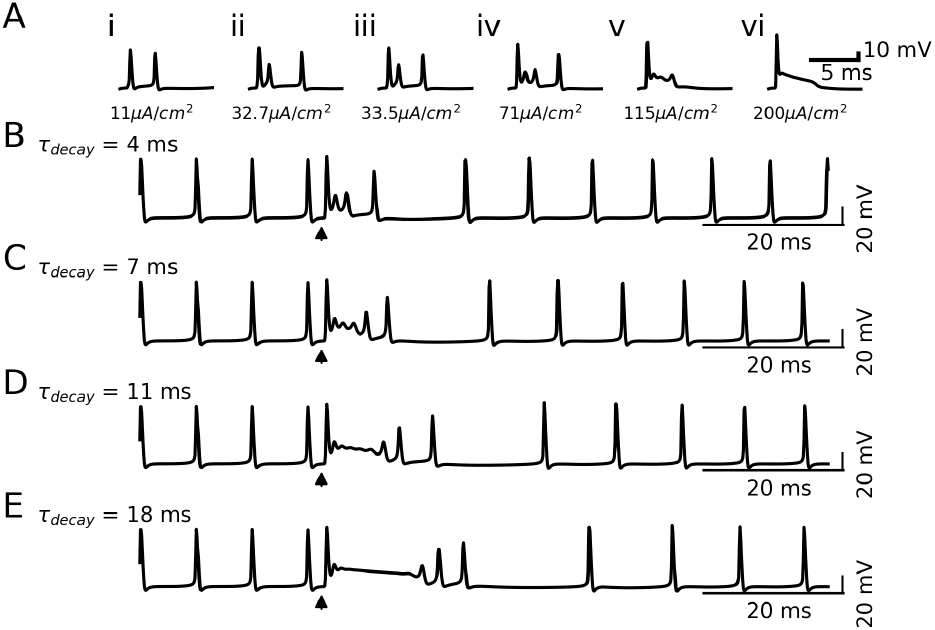
Response of the five-current model to input is similar to the response of the three-current model. Simulated complex spike waveforms in the five-current model with added calcium and SK channels behave similarly to increasing time constants describing the decay of the injected climbing fibre input as the three-current model. Complex spike waveform simulations when increasing *τ*_−_ from 4 ms **(A)**, 7 ms **(B)**, 11 ms **(C)** and 18 ms **(D)**. Arrows indicate the times of the simlulated climbing-fibre-like current injections.

Changes in the activity of the SK channel was found to influence Purkinje cell simple spike rate. Increases in the maximum conductance through either the calcium-activated potassium channel or the calcium channel prolonged the duration of the post-complex spike pause in simple spike activity and reduced baseline simple spike rates, whereas decreases in maximal conductance reduced the duration of the pause and increased simple spike firing frequency, see Fig. 7. Increasing 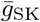 in the model caused a slight increase in simple spike rate above baseline levels after the pause. This is reminiscent of simple spike modulation observed in Purkinje cells in vivo (Burroughs et al., 2017).

**Figure 7:**
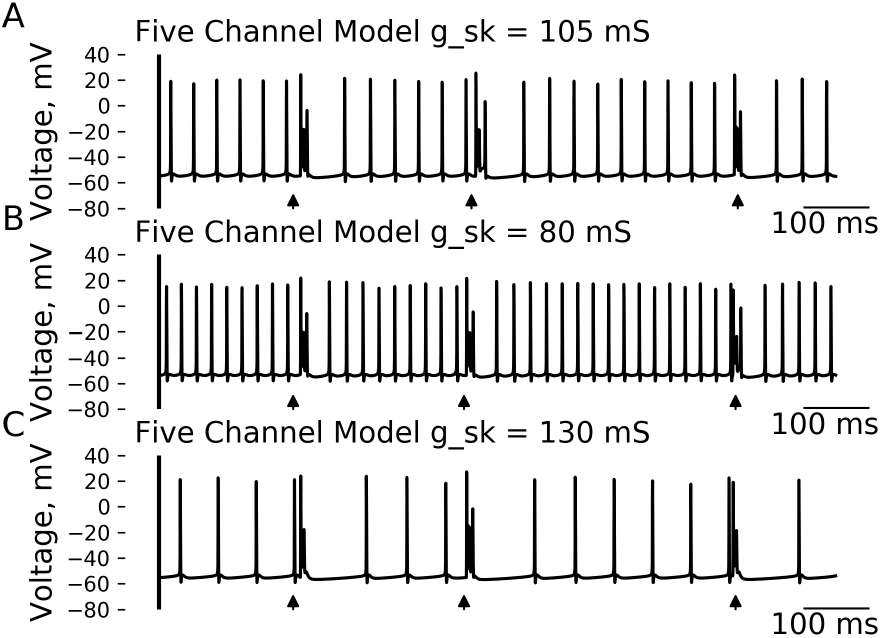
The model is robust to changes in parameter values. Changing the parameter 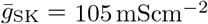 **(A)** influences simple spike rate and pause duration. Decreasing 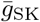 from 105 mScm^-2^ to 80 mScm^-2^ leads to an increase in background simple spike rate and a reduction in the duration of the post-complex spike pause in simple spike activity **(B)**, whereas increasing 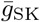 from 105 mScm ^2^ to 130 mScm^-2^ decreases simple spike rate and increases pause duration **(C)**. Arrows indicate the time of climbing-fibre-like current injection. The model was robust to small changes in other parameter values.

## 4 Discussion

The present study tested, through computational modelling, the possibility that key features of Purkinje cell complex spike waveform can be generated by a limited number of channel-dynamics. Our three-current model exhibits the main features of the complex spike: a high-amplitude initial spike that is followed by a succession of smaller amplitude spikelets with the first spikelet having a smaller amplitude than those that follow. In the five-current model, which extends the three-current model by including a P/Q type calcium channel and a calcium-activated potassium channel, some of the interactions between complex spike activity and background simple spike behaviour are also observed in simulations.

In these models, the dynamics of the resurgent sodium channel has a crucial role in producing the complex spike waveform, which is in line with experiments (Raman and Bean, 1997; Nolan et al., 2003; Khaliq and Raman, 2006). After the initial spike of the complex spike, channels become sequestered in the open-but-blocked state so that the following spikelets are stunted relative to the initial spike and blocked channels returning to the open state help sustain the depolarization for subsequent spikelet generation.

In fact, somatic complex spikes are often thought to be generated by an interplay between the resurgent sodium current (Raman and Bean, 1997, 2001; Khaliq et al., 2003; Khaliq and Raman, 2006) and the current through the fast potassium channel K_v_3.3 (Zagha et al., 2008; Hurlock et al., 2008; Veys et al., 2013); this is the point-of-view developed in the two models presented in this paper and these models are intended as a demonstration that this is a plausible description of complex spike generation.

In Purkinje cells, resurgent sodium current is conducted through Na_v_ 1.6 channels (Raman and Bean, 1997) and is estimated to contribute 40% of total sodium current generated during a simple spike (Raman and Bean, 2001), although others suggest this figure is only 15% (Levin et al., 2006). Resurgent sodium channels are found in numerous neuronal types throughout the brain and spinal cord (Osorio et al., 2010), and are unusual in that once they open they either inactivate or enter into a reversible open-but-blocked state that is non-conducting, see Table 1. Phosphorylation of the cytosolic *β_4_* subunit terminal may be responsible for this form of open channel block (Grieco et al., 2002). Upon repolarisation, channels in the open-but-blocked configuration return through the open configuration to either a closed or inactive state.

In Purkinje cells, the K_v_3.3 potassium channel is responsible for rapid repolarisation (Veys et al., 2013) because of its fast channel kinetics (Rudy et al., 1999; Rudy and McBain, 2001). These fast potassium dynamics enable spikelet frequency to reach their observed ~ 600 Hz frequency (Warnaar et al., 2015; Burroughs et al., 2017). Somatic, but not dendritic, K_v_3.3 channels are critical to repetitive spikelet generation and define the shape of the Purkinje cell complex spike (Hurlock et al., 2008; Zagha et al., 2008; Veys et al., 2013).

The multi-compartmental model developed in De Schutter and Bower (1994a,b,c) also shows a complex spike waveform under some conditions. These complex spikes do not have the architecture of relative spikelet amplitudes and inter-spikelet intervals subsequently observed *in vitro* and the spikelets are generated by a different mechanism: the interplay of a delayed rectifier potassium current and the fast, transient sodium current. The multi-compartmental model has a delayed rectifier potassium current with unusually large opening and closing rates, exceeding experimental values by a factor of five (Yamada et al., 1989); this plays a similar role to the Kv3.3 fast delayed rectifier potassium gate, (Veys et al., 2013; Zagha et al., 2008) used in the models described here: Kv3.3 is necessary for the rapid repolarisation of the membrane following each spike or spikelet, as noted experimentally in Zagha et al. (2008); Veys et al. (2013). Zang et al. (2018) have recently developed a full scale compartmental model, this model does simulate the complex spike, but only in the contex of a very large model which does not clarify which aspect of the model are required for complex spike generation.

In the models presented here an unusually large leak conductance of 2 mScm^-2^ was necessary to accurately represent the dynamics of complex spikes recorded *in vitro*, namely the high spikelet firing frequency, the latencies between individual spikelets and the progressive increase in spikelet amplitude throughout the complex spike; thus, the model suggests that the passive resistance of the Purkinje cell somatic membrane is low. Evidence in support of this has been found experimentally, as a disparity between somatic and dendritic membrane resistance is present in Purkinje cells (Rapp et al., 1994). This disparity is also a feature of the model in De Schutter and Bower (1994a,b,c) where somatic membrane resistance was 440 Ω cm^2^, which is equivalent to a conductance of ≈ 2.3mScm^-2^, compared to 11kΩcm^2^ in the dendrites. Membrane resistance determines the membrane time constant which, in turn, determines the time scale over which the cell responds to input. With low resistivity, the Purkinje cell soma will have a smaller membrane time constant than the dendrites, facilitating the generation of very fast spikelets (Warnaar et al., 2015).

The shape of the complex spike waveforms, including complex spike duration, spikelet amplitudes and spikelet timing, generated in the two models is robust to changes in parameter values. Both fixed and free parameters can be changed without significantly altering these complex spike dynamics. The exception to this is the applied background current: as well as reducing simple spike activity, reducing the background current increases the amplitude of spikelets.

Calcium-activated potassium channels are included in the five-current model because they contribute to interactions between complex spikes and simple spike activity and regulate Purkinje cell output (Tank et al., 1998; McKay et al., 2007). SK channels play an important role in limiting the firing rate of Purkinje cell simple spike activity (Womack and Khodakhah, 2003; Egorova et al., 2014), which is often correlated with complex spike activity. In the absence of complex spiking, simple spike firing frequencies increase to exceptionally high levels (Cerminara and Rawson, 2004). SK channels are also responsible for afterhyperpolarisations that follow bursts of action potentials (Ohtsuki et al., 2012), which may underlie the climbing fibre-induced pause and downregulation of simple spikes (Jin et al., 2017). The model of the calcium-activated potassium current used here and taken from Gillies and Willshaw (2006) is one of a number of models for the SK channel, for example, the alternative model used in Griffith et al. (2016) based on Chay and Keizer (1983); Xia et al. (1998) has a steeper and quicker modulation of the channel by calcium concentration. Furthermore, the model from Gillies and Willshaw (2006) simplifies the original dynamics described in Hirschberg et al. (1998) and a future elaboration of the Purkinje cell model presented here might consider how the richer temporal dynamics in Hirschberg et al. (1998), with different calcium concentration decay rates, might affect the behaviour of the model.

The dynamics of intracellular calcium concentration is based on Eilers et al. (1995) where it was demonstrated that intracellular calcium signalling within the Purkinje cell soma is not temporally nor spatially uniform. Within a narrow submembrane shell of 2 to 3 *μ* m, calcium transients are large and fast when compared to the interior of the soma. The calcium concentration within this narrow submembrane region was modelled in the present study. Intracellular calcium dynamics, in particular the time course of calcium concentration decay, was found to modulate simulated Purkinje cell simple spike activity. However, it should be noted that the five-current model includes parameters, such as the submembrane shell thickness and decay in calcium concentration, that are not well-specified experimentally and in any case likely depend on somatic morphology, something that is variable across Purkinje cells.

Although the three-current and five-current models differ in how the complex spike affects simple spiking, there is almost no difference in the complex spike waveform in the two models, Fig. 6. On the other hand, the shape of the pre-synaptic climbing fibre input seems to be critical in shaping complex spike waveforms and so changes in this input provides a possible mechanism for their variation.

Purkinje cells *in vitro* sometimes show a bimodal pattern of simple spike activity, cycling between an *up* and a *down* state. The up state is characterised by high simple spike rates whereas in the down state simple spiking is quiescent. Several models of Purkinje cell activity have attempted to describe this bimodality (Forrest, 2014; Forrest et al., 2012; Llinas and Sugimori, 1980b; Loewenstein et al., 2005; McKay et al., 2007; Williams et al., 2002). Combined theoretical and experimental work has shown that Purkinje cells exhibit a phenomenon known as inverse stochastic resonance. This means they can efficiently transitions between different functional regimes depending on the variability of their synaptic input (Buchin et al., 2016). The models presented in the current study do not exhibit bimodal behaviour and the bimodality is not generally representative of the firing patterns observed *in vivo* (Cerminara and Rawson, 2004; McKay et al., 2007; Schonewille et al., 2006).

The simplified Purkinje cell models presented here do not include some of the channels known to be expressed in the Purkinje cell membrane. These include the HCN1 channel, which may be responsible for bi-modal patterns of activity (Loewenstein et al., 2005), the calcium-activated intermediate conductance (IK) and big conducance (BK) potassium channels, which contribute to the regulation of Purkinje cell excitability and rhythmicity (Cheron et al., 2009), the persistent sodium channel, which is involved in nonlinear synaptic gain, plateau potentials and subthreshold oscillations (Kay et al., 1998), the K_v_1.1 channel, which prevents hyperexcitability (Zhang et al., 1999), the T-type calcium channels, which play a role in rhythmicity and bursting behaviour, for review see Cain and Snutch (2010), and the inwardly rectifying potassium channels, which are activated by GABA B receptors (Tabata et al., 2005). These currents are likely to play a role in modulating Purkinje cell activity.

The models presented in this study are tuned to match complex spikes recorded *in vitro* in Davie et al. (2008). This example is chosen because it typifies features of the complex spike waveform observed in numerous *in vitro* studies (Zagha et al., 2008; Zhang et al., 2017; Monsivais et al., 2005; Khaliq and Raman, 2005) and in an *in vivo* study, Warnaar et al. (2015): *in vivo* studies that give wave form data are less common. These feature, which are broadly stereotypical are

1. the first spike is similar in shape to the simple spike,
2. the first spikelet is smaller in amplitude than subsequent spikelets,
3. spikelets get further apart with increasing current injections or spikelet number.

The comples spikes recorded in Davie et al. (2008) were also used as a target beause the complex spike waveforms were recorded in response to increasing injections of climbing fibre like currents. This describes the complex spike waveform response to different somatic inputs and provides a useful range of responses to use as a target for model fitting. The typical features of complex spikes and the response to varying input are exemplified by the fitted model. The five-current model also captures the extended refractory period that follows a complex spike; since this feature is easily seen in extra-cellular recordings, it is supported by a larger range of *in vivo* data, see for example Granit and Phillips (1956); Latham and Paul (1970); Bloedel and Roberts (1971); Armstrong and Rawson (1979); Bauswein et al. (1983); Sato et al. (1992); Burroughs et al. (2017).

However, complex spikes do exhibit large amounts of variability (Burroughs et al., 2017; Warnaar et al., 2015). Interestingly both the three- and five-current models presented here can be used to describe these variations: for example, the difference in complex spikes seen in Monsivais et al. (2005) matches the variation in waveform seen in Fig. 6.

Obviously, restricting the model to a single somatic compartment is a substantial simplification. Along with removing any affect of the complex somatic morphology, it fails to model the coupling between the soma and dendrite. This is significant because the extensive dendritic arborisations that characterise Purkinje cells means that there are space clamp limitations associated with recording from Purkinje cells. Removing the coupling to the dendrite is an approximation and rules out the possibility, in our models, that current flow from the soma to the dendrite might have a role in the production of complex spikes. However, as discussed above, the complex spike has been shown to be generated only within the proximal axon, the structure of the dendritic input is lost during transmission to the soma and there is a large discrepancy between the membrane resistance in the soma with current from the soma to the dendrite likely to be small. As such, since one-compartment models are considerably easier to interpret, this suggests a one-compartment model is useful at this level of approximation.

Experimentally, a positive correlation has been observed between simple spike rate and the number of spikelets making up a subsequent complex spike waveform *in vivo* (Burroughs et al., 2017). While the five-current model did simulate the extended refractory period that follows a complex spike, it was unable to capture these more complicated interactions between complex spike waveform changes and simple spike dynamics, suggesting that these may result from mechanisms located outside of the soma, such as dendritic computations, synaptic plasticity or network effects. In the future these models will be useful as a component in a larger model of local cerebellar circuitry.

Here the model can capture a more physiological response to experimentally recorded trains of parallel fibre and climbing fibre activity and be useful in identifying how the network responds to complex spike waveform and simple spike activity. The models presented here could be used to address a number of further questions including, but not limited to, the influence of synaptic plasticity at single synapses on Purkinje cell activity, the temporal significance of spike activity patterns, the contribution of individual channels to specific electrophysiological properties and how these are modulated by neurotransmission and intracellular pathways. This will enable us to gain a better insight into the electrodynamics supporting cerebellar function and to make predictions about network dynamics, including those that occur during learning or aberrant activity associated with disease, which could then be tested experimentally.

## Acknowledgements

Special thanks go to Beverley Clark and Jennifer Davie for providing us with *in vitro* complex spike data to which this model is tuned.

## Code Availability Statement

The Python code used to create the figures is available at github.com/PurkinjeCellSimpleModel.

## Footnotes

1 `scipy.optimize.minimize`

